# Reading-out task variables as a low-dimensional reconstruction of neural spike trains in single trials

**DOI:** 10.1101/643668

**Authors:** Veronika Koren, Ariana R. Andrei, Ming Hu, Valentin Dragoi, Klaus Obermayer

## Abstract

We propose a new model of the read-out of spike trains that exploits the multivariate structure of responses of neural ensembles. Assuming the point of view of a read-out neuron that receives synaptic inputs from a population of projecting neurons, synaptic inputs are weighted with a heterogeneous set of weights. We propose that synaptic weights reflect the role of each neuron within the population for the computational task that the network has to solve. In our case, the computational task is discrimination of binary classes of stimuli, and weights are such as to maximize the discrimination capacity of the network. We compute synaptic weights as the feature weights of an optimal linear classifier. Once weights have been learned, they weight spike trains and allow to compute the post-synaptic current that modulates the spiking probability of the read-out unit in real time. We apply the model on parallel spike trains from V1 and V4 areas in the behaving monkey *macaca mulatta*, while the animal is engaged in a visual discrimination task with binary classes of stimuli. The read-out of spike trains with our model allows to discriminate the two classes of stimuli, while population PSTH entirely fails to do so. Splitting neurons in two subpopulations according to the sign of the weight, we show that population signals of the two functional subnetworks are negatively correlated. Disentangling the superficial, the middle and the deep layer of the cortex, we show that in both V1 and V4, superficial layers are the most important in discriminating binary classes of stimuli.

## Introduction

A half century ago, pioneers of neuroscience have stated the following: “At present we have no direct evidence on how the cortex transforms the incoming visual information. Ideally, one should determine the properties of a cortical cell, and then examine one by one the receptive fields of all the afferents projecting upon that cell.” (Hubel and Wiesel, 1962, Journal of Physiology, [1]). While lots of insights in the computations in cortical circuits have been made in the meantime, the question, posed by Hubel and Wiesel, has not yet found a clear answer [2]. Addressing this question requires observing the activity of many neurons simultaneously and has demanded an important progress in recording techniques. Besides advances on the experimental side, a major challenge is to interpret these rich datasets and to understanding the underlying principles of cortical computation [3, 4]. One of the biggest conceptual gaps to bridge is between sensory processing and animal’s behavior, addressed by decision-making studies [5–8]. Linking behavioral choices with the neural activity in the sensory areas requires the understanding of the transformation between sensory and decision-related signals. On the one hand, spiking patterns of neural populations in sensory areas are highly variable across trials [9], and can be described as a probabilistic process. The choice behavior of animal agents, on the other hand, appears to be highly precise and coherent with respect to the incoming (natural) stimuli. Even though the behavior is noisy [10], and prone to errors due to wrong internal representation [11], this might be so because it can adapt to perturbations and the uncertainty in the environment [12]. Our main question here is how does the brain transform a high-dimensional probabilistic signal, enacted by spike trains of cortical populations, into a reliable signal, that presumably underlies coherent animal behavior.

Recent theoretical and modeling work has shown that it is possible to read-out a deterministic population signal from variable spike trains of a spiking neural network [13]. In [13], it is assumed that the population signal is encoded at the network level and cannot be accounted for by the observation of single neurons. The population signal is distributed among single neurons in a non-linear fashion, giving rise to a distributed code. The distributed code maps from the high-dimensional space of spike trains of many neurons to the low-dimensional space of the population signal. If the coding function of individual neurons is redundant within the network, many different spiking patterns can be decoded as the same population signal. Such a coding scheme therefore allows to reconcile variable spike trains with a deterministic signal, that might underlie animal’s behavior. While efforts have been made to design an efficient network that is biologically plausible [14], no convincing evidence for such a computation in biological ensembles has been presented so far.

In the present study, we apply theoretical propositions of the model with the distributed code [13, 15] to experimental data. Our goal is to connect the theory on representation of an abstract and arbitrary population signal to behaviorally relevant variables in the biological brain. Aforementioned studies suggested that the activity of a spiking network can be decoded by weighting spikes ([13], [15], [16], [17]), where decoding consists in transforming spike trains of many neurons, a high-dimensional variable, into a low-dimensional population signal. The core of the transformation is to weight the spike trains of individual neurons by their decoding weight and sum across neurons, giving a low-dimensional representation of network’s spiking activity. Since these studies show the proof of a concept, they utilize random weights. Here, we propose that weights depend on the computational task at hand, in our case, discrimination of binary stimulus classes. We also propose that in the biological setting, such a low-dimensional signal corresponds to the synaptic current received by a read-out neuron, modulating its probability of spiking.

Our aim is to build a framework that is a minimal but generic model of stimulus processing in the brain. In general, decoding models either do *classification* or *stimulus reconstruction* [18]. Here, we break the decoding problem in two parts, 1) learning of weights and 2) low-dimensional reconstruction of parallel spike trains. We assume that learning of weights occurs through supervised learning and is a *classification* problem. Once the weights are learned, they are fixed and we use them for *stimulus reconstruction* in real time. Methods of supervised learning have the advantage of dealing naturally with multivariate signals and of insuring that the model generalizes on yet unseen data, but often suffer from lack of interpretability of results [19]. We combine *classification* with *reconstruction* to design a read-out method that is interpretable in a biological setting. While our method cannot establish a causal relationship between neural activity and behavior (see [20]), it does compute the upper bound of the information about the choice variable that can be extracted from neural activity of observed neural ensembles [19]. Whether optimal weights are plausible in a biological neural network is an open question [18]. We address this issue by showing that optimality of weights can be relaxed, with only a small loss of decoding power. To truly remove the danger of a “black-box” methodology, we also explicitly demonstrate the role of each source of information for the model by removing a specific type of information and evaluating the effect of such a perturbation on the read-out.

In short, this study attempts to bridge the gap between abstract models of computation in neural networks and activity of neural populations, recorded *in vivo* in visual areas V1 and V4. It addresses decoding of multivariate signals that pertain to correct choice behavior and tries to bring insights about “...how do the connectivity and dynamics of distributed neural circuits give rise to specific behaviors and computations” (Gao & Ganguli, 2015, *Curr. Op. in Neurobiology*, [3]).

## Materials and methods

### Ethics statement

All experiments performed in this study were conducted in accordance with protocols approved by The Animal Welfare Committee (AWC) and the Institutional Animal Care and Use Committee (IACUC) for McGovern Medical School at The University of Texas Health Science Center at Houston (UTHealth), and met or exceeded the standards proposed by the National Institutes of Health’s Guide for the Care and Use of Laboratory Animals.

### Animal subjects

Two male rhesus macaques (Macaca mulatta; M1, 7 years old, 15kg; M2, 11 years old, 13kg) were used in this study. Subjects were housed individually (after failed attempts to pair house) in cages sized 73 x 69 x 31 or 73 x 34.5 x31 inches, in close proximity to monkeys in adjacent cages, allowing for visual, olfactory and auditory contact. Toys were given in rotation, along with various puzzles, movies and radio programming as environmental enrichments. Monkeys were fed a standard monkey biscuit diet (LabDiet), that was supplemented daily with a variety of fruits and vegetables. Subjects had been previously trained to perform visual discrimination task, and each implanted with a titanium head post device and two 19mm recording chambers (Crist Instruments) over V1 and V4. All surgeries were performed aseptically, under general anesthesia maintained and monitored by the veterinary staff from the Center for Laboratory Animal Medicine and Care (CLAMC), with appropriate analgesics as directed by the specialized non-human primate veterinarian at CLAMC. During the study the animals had unrestricted access to fluid, except on days when behavioral tasks were performed. These days, animals had unlimited access to fluid during the behavioral task, receiving fluid for each correctly completed trial. Following the behavioral task, animals were returned to their home cage and were given additional access to fluid. The minimal daily fluid allotment was 50ml/kg (monkeys were weighed weekly), though monkeys could drink more through their participation in the task. During the study, the animals’ health and welfare was monitored daily by the veterinarians and the animal facility staff at CLAMC and the lab’s scientists, all specialized with working with non-human primates.

#### Experimental setup

Animals performed a visual, delayed-match-to-sample task. The trial started after 300 ms of successful fixation within the fixation area consisted in displaying the target and the test stimuli, naturalistic images in black and white, with a delay period in between. The target and the test stimuli were either identical (condition “match”) or else the test stimulus was rotated with respect to the target stimulus (condition “non-match”). The target and the test stimuli were shown for 300 ms each while the delay period had a random duration between 800 and 1000 ms. Random duration of the delay period prevented that the subject anticipates the time of arrival of the second stimulus. Visual stimuli were images in black and white and represented an outdoor scene. The identity of the stimulus changed on every trial, but always fell into one of the two categories, “match” and “non-match”, where “match” indicates an identical pair of target and test stimuli, and “non-match” indicates the rotation of the test stimulus with respect to the target. The task of the animal was to decide about the similarity of the target and the test stimuli by holding a bar for “different” and releasing the bar for “same”. The subject was required to respond within 200 and 1200 ms from the time the test stimulus was off, otherwise the trial was discarded. The difference in orientation of the test stimulus ranged between 3 and 10 degrees and was calibrated on-line in order to have on average 70 percent correct responses on non-matching stimuli. The subject was rewarded for a correct response with fruit juice.

Recording were made with laminar electrodes with 16 recording channels. In part of sessions, recordings were made in V1 and V4 simultaneously, with one laminar electrode in each area, while in other sessions, only V1 has been recorded. The position of the electrode was calibrated in such a way that neurons from the two areas had overlapping receptive fields. The multi-unit signal and the local field potential were recorded in 20 recording sessions in V1 and in 10 recording sessions in V4. We analyzed the activity of cells that responded to the stimulus by increasing their firing rate at least four-fold with respect to their baseline. We used the activity of all neurons that obeyed that criterion, which gave 160 neurons in V1 and 102 neurons in V4. The number of trials was roughly balanced across conditions (tab.1).

**Table 1.** Number of trials per condition.

| Area | Condition | average nb. trials | Area | Condition | average nb. trials |
| --- | --- | --- | --- | --- | --- |
| V1 | “match” | 118 | V4 | “match” | 99 |
|  | “non-match” | 107 |  | “non-match” | 76 |

#### The spike train and the population PSTH

The following analyses were done with Matlab, Mathworks, version R2017b.

The spike train of a single neuron *n* in trial *j* is a vector of zeros and ones,

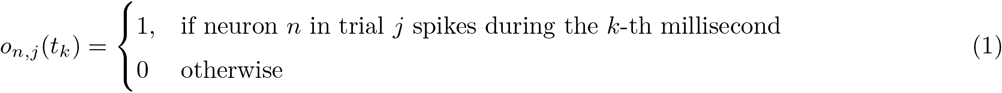

where *n* = 1, …, *N* is the neural index, *j* = 1,.., *J* is the trial index and *k* = 1*, …, K* is the time index with step of 1 millisecond. The population PSTH is computed by averaging spike trains across neurons and across trials.

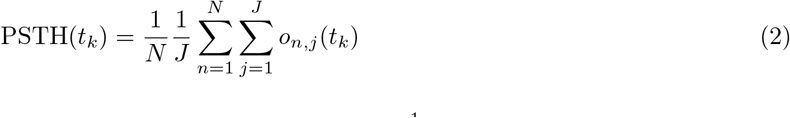

The PSTH is convolved with a Gaussian kernel, 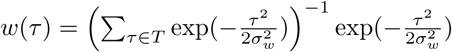 with variance 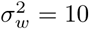ms and support T = {−10, …, 10} ms.

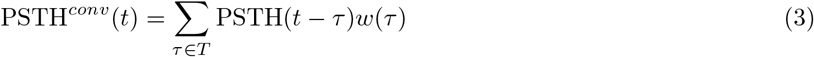

#### Estimation of decoding weights

Trials were split into training and validation set, with *j* = 1, …, *J′* trial indexes for the training set and *j* = *J′* + 1, …, *J* trial indexes for the validation set. We used the training set to compute decoding weights and the validations set to apply weights to spike trains and compute the population signal. The training and the validation set were non-overlapping and utilized half of the available trials each. The split into training and validation set was cross-validated with the Monte Carlo method. In every cross-validation run, the trial index is randomly permuted, without repetition, and the data is split into training and validation set. Throughout the paper, we used 100 cross-validations and the reported results are averages across cross-validations. All computations are done for each recording session independently.

##### Constructing features

Cortical neurons adjust the strength of their synapses through learning. Here, we assume that after learning, synaptic weights reflect the role of each neuron within the population for the computational task that the network has to solve, which is binary classification of stimulus classes. Since the animal is rewarded in correct trials, we assume that the reward signals enacts a teaching signal, formalized in the setting of supervised learning. Moreover, a single cortical neuron typically receives synaptic inputs from many projecting cells [9]. For this reason, we assume that learning of synaptic weights relies on the joint activity of simultaneously recorded neurons, where interactions between neurons are taken into account.

Decoding weights were computed as the feature weights of the linear Support Vector Machine (SVM, [21]). The latter has been chosen for its optimality and for the interpretability of results in the biological setting. The SVM was trained on the spike count statistics. The spike count of the neuron *n* in trial *j*, 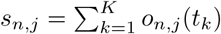, is computed in target and test time windows, corresponding to the interval of [0, 400] milliseconds with respect to the onset of the target and the test stimuli. Rather than the absolute spike count, we assume that the relevant signal for learning is the deviation of the spike count from the baseline. Spike counts are therefore *z*−scored, for each neuron independently,

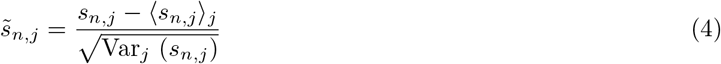

where 〈*s*_*n,j*_〉_*j*_ is the empirical mean and Var_*j*_ (*s*_*n,j*_) is the empirical variance across trials.

##### Model fitting and extraction of weights

Let’s have an N-dimensional vector of activities in trial *j*, 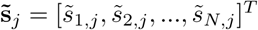, and the class label *y*_*j*_ ∈ {−1, 1}, where *y* = −1 is the label for condition “non-match” and *y* = 1 is the label for condition “match”. Linear SVM searches for an *N* − 1-dimensional plane (hyperplane) that optimally separates points in conditions “match” and “non-match”,

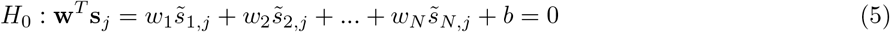

where **w** = [*w*_1_, …, *w*_*N*_]^*T*^ is a *N* -dimensional vector of feature weights and *b* is the offset of the hyperplane from the origin. Given data points, the hyperplane *H*_0_ is fully described by the vector of feature weights and the offset, where the vector of feature weights determines its direction (i.e., **w** is perpendicular to *H*_0_).

The optimization problem that the linear SVM solves is, in its primal form, expressed with a Lagrangian,

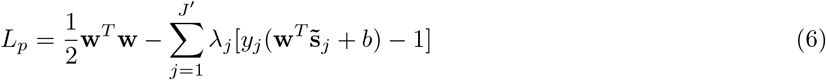

with *λ*_*j*_ ≥ 0 ∀*j* the Lagrange multiplier. The minimization of the Lagrangian results in Lagrange multipliers equal to zero for as many samples as possible, and only a subset of samples, those that lie on the margin, will be used to determine the separation boundary. Those samples are called the support vectors: **v**_*q*_ = [*v*_1,*q*_, *v*_2,*q*_, …, *v*_*N,q*_], *q* = 1, …, *Q*, *v*_*n,q*_ ∈ ℝ. Minimizing the Lagrangian, i.e., differentiating the Lagrangian with respect to the weight vector **w** and setting the derivative to zero, one obtains the expression for the weight vector.

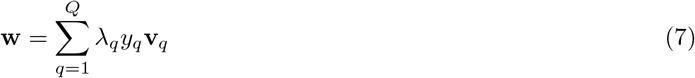

We normalize the weight vector with the *L*^2^ norm,

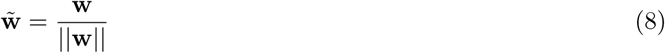

with 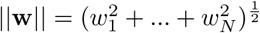. The normalized weight vector, 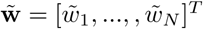, was computed in every recording session (i.e., for simultaneously recorded neurons). It associates the activity of each neuron within the population with neuron’s function for classification. Since the classification is done in the high-dimensional space, the weight of a single neuron is relative to activities of other neurons from the population, and the weight vector naturally takes into account inter-neuron interactions.

In general, it is not necessarily possible to linearly separate all the input samples, not even in the training data. The classification model of the SVM optimizes the separating hyperplane also with respect to the data points that lie on the wrong side of the margin (slack points). The regularization parameter determines how much do slack points contribute to the error that the model is minimizing. If slack points contribute strongly to the error, the model converges to a narrow margin, which might result in bad generalization of the hyperplane to the test data (overfitting). Here, we chose the regularization parameter with 5-fold cross-validation on the training set. The training set was split into 5 folds, the classifier was trained on 4 folds and validated on the remaining fold. Each validation sample is classified as the true positive, true, negative, false positive or false negative.

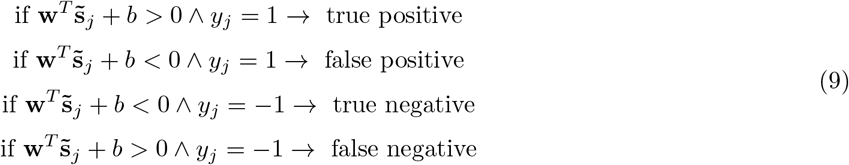

The performance of the classifier is evaluated with balanced accuracy,

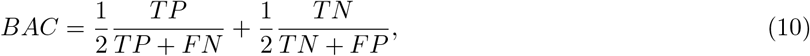

where *TP*, *TN*, *FP* and *FN* are the number of true positive, true negative, false positive and false negative samples, respectively. Balanced accuracy accounts for imbalanced classes, that is, different number of trials in conditions “match” and “non-match”. When the 5 combinations of training/validation folds are exhausted, we compute the average balanced accuracy across folds. Iterating this procedure for a range of regularization parameters, *C* ∈ {0.0012, 0.0015, 0.002, 0.005, 0.01, 0.05, 0.1, 0.5}, we chose the regularization parameter that maximized the balanced accuracy. The regularization parameter is fitted to every recording session independently.

Notice that the range of the weight vector (eq. 7) depends on the regularization parameter *C*, and is therefore different across recording sessions. Since we combine results from many recording sessions, normalization of the weight vector (eq. 8) is necessary to keep decoding weights in the same range across recording sessions [18].

#### Low-dimensional population signal

##### The population signal as the weighted sum of spikes

Imagine a population of N neurons that project to a read-out neuron. Every spike of a projecting neuron creates a small jump in the membrane potential of the read-out neuron, followed by a decay towards the baseline [22]. Moreover, spikes of all projecting neurons are summed up in the membrane potential of the read-out neuron [22]. Consider the spike train of *N* simultaneously recorded neurons in trial *j* and in time step *t*_*k*_,

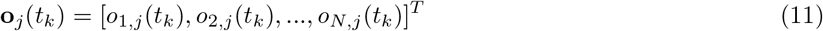

where *j* = *J′* + 1, …, …, *J* are trials from the test set. Trials in condition “non-match”, *j* = *J′* + 1, …, *J′′*, are followed by trials in condition “match”, *j* = *J′′* + 1, …, *J*. The transformation of spike trains into a low-dimensional population signal consists in multiplying the spike train of each neuron with the corresponding weight, summing across neurons and convolving with an exponential kernel. Equivalently, this can be written as the projection of spike trains on the vector of decoding weights,

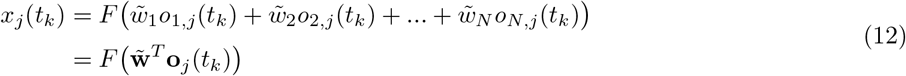

where *F* (*y*) is the transfer function. The transfer function is defined as the convolution with an exponential kernel,

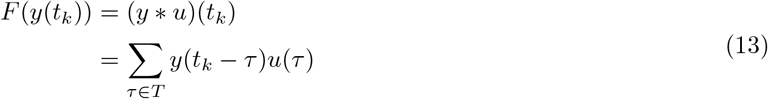

with *u*(*t*) = *exp*(−*λt*). Convolution with an exponential kernel models the causal effect of the presynaptic spike on the neural membrane of the read-out neuron. Note that the transformation applies to a specific trial and maintains the temporal dimension of the spike train. The only manipulation is to reduce the dimensionality from N (dimensionality of the spike train) to 1 (dimensionality of the population signal). We compute the deviation of the resulting signal from the mean,

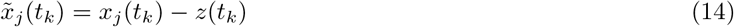

where *z*(*t*_*k*_) is the average population signal across trials from both conditions.

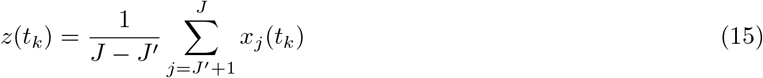

The low-dimensional signal is then averaged across trials, distinguishing trials from condition “match” and “non-match”.

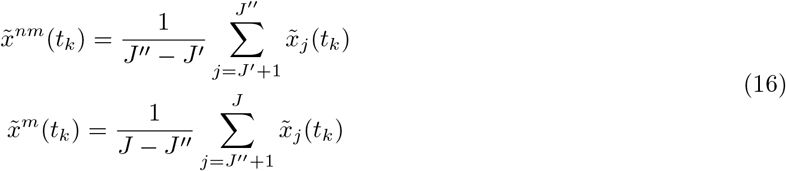

The significance of the discrimination between the population signal in “match” and “non-match” is evaluated with he permutation test. The test statistics is the difference of population signals.

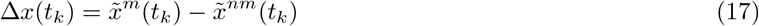

We compare ∆*x*(*t*_*k*_) with 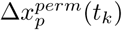, where the latter has been computed with random permutation of class labels for “match” and “non-match. The permutation is done in the training as well as in the validation step. In the training step, decoding weights are computed with SVMs that are trained on randomly permuted class labels. In the validation step, we randomly permute class labels of parallel spike trains, keeping the same label across neurons. The whole procedure is repeated *nperm*-times and gives a distribution of results for each time step. When the result of the true model appears outside of the distribution of results for the null model, the difference of signals in conditions “match” and “non-match is considered to be significant.

##### The population signal of neurons with positive and negative weights

We separate the population of simultaneously recorded neurons with respect to the sign of the decoding weight, distinguishing neurons with positive weight (*plus* neurons) and negative weight (*minus* neurons). Weight vector of *plus* 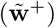 and *minus* neurons 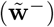, is defined by replacing weights of the opposite sign with zero. This way, spikes of neurons with the opposite sign are weighted by a zero weight and do not contribute to the projection. The population signal is computed with each of the two weight vectors,

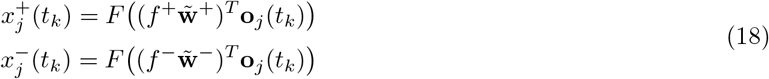

where *f* ^+^ (*f* ^−^) stands for the correction factor for the number of *plus* (*minus*) neurons. The correction factor is computed as 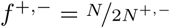, where *N* ^+^ (*N* ^−^) is the number of *plus* (*minus*) neurons. We have, on average, 58 % of *plus* neurons in V1 and 54 % in V4 during target, and 54 % of *plus* neurons in V1 and 62 % in V4 during test. To be able to compare results across the two neuronal types, we scale the weight vector with the correction factor.

We compute the deviation of the population signal from the mean,

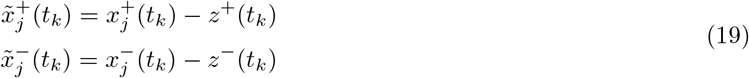

with *z*^+^(*t*_*k*_) and *z*^−^(*t*_*k*_) are the sign-specific average of population signals.

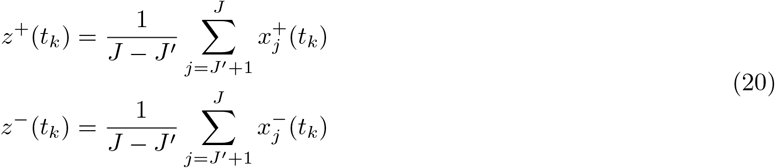

Finally, we average each of the signals across trials, distinguishing conditions “match” and “non-match”.

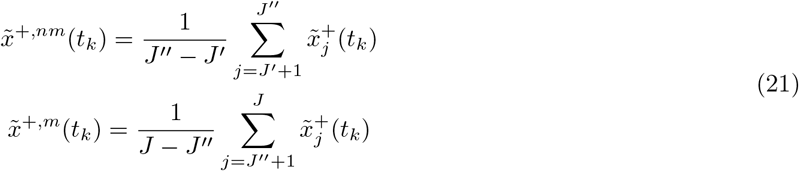

Same follows for *minus* neurons.

The significance is evaluated with the permutation test. The test statistic is the sign-specific difference of signals in conditions “match” and “non-match”, e.g., for *plus* neurons:

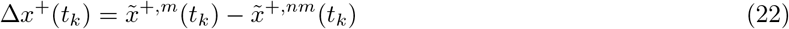

The null model is computed with the random permutation of class labels when training the classification model as well as in the reconstruction step. In addition, we use a random assignment to the class of *plus* and *minus* neurons by randomly permuting neural indexes before splitting neurons in *plus* and *minus* neurons.

##### The population signal in cortical layers

We distinguish three cortical layers, superficial (supragranular, SG), middle (granular, G) and deep layer (infragranular, IG, [23]). The method for determining cortical layers is described in the last section of methods, “Determining cortical layers from the current source density”. We define three layer-specific weight vectors, 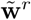, *r* ∈ {*SG, G, IG*}. The layer-specific weight vectors take into account weights of neurons from the specific layer, while weights of other neurons are replaced with zero. The reconstruction in layer *r* is defined as follows:

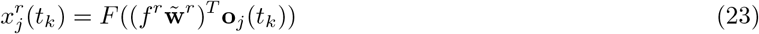

where 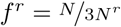 is the correction factor for the number of neurons across layers, with *N*^*r*^ the number of neurons in layer *r*. For the rest, the computation of layer-specific signals is identical as for the *plus* and *minus* neurons, substituting sign-specific with layer-specific signals.

##### Disentangling sources of information for discrimination of conditions “match” and “non-match”

The population signal contains several sources of information that might inform successful discrimination of conditions “’match” from “non-match”. Useful information could come from the weight vector and/or from the spike timing (eq. 12). Note that the weight vector is computed utilizing spike counts and therefore does not contain the information on spike timing. To investigate the contribution of different sources of information, we remove a particular source while keeping others intact, and test how such a perturbation affects the discrimination. If a particular source of information is critical, its removal will compromise the discrimination. The information is removed by replacing the true statistics with random statistics.

We remove the information in the weight vector by substituting the true weight vector with a random vector. The random vector is given by drawing N random samples (where N is the number of neurons in the recording session) from the uniform distribution with the same range as the range of the regular weight vector. We compute the population signal (eq. 12 - 21) utilizing true spike trains and the random weight vector. The information in decoding weights can be further split into the information contained in the sign of weights (positive or negative) and in the modulus of weights (the absolute value of each element of the weight vector). The information contained in the sign of weights is removed by drawing N random samples from the uniform distribution, collecting their signs and applying them to the regular weight vector. The information in the modulus is removed by replacing the modulus of each element of the regular weight vector with the modulus of a random vector, but keeping the correct sign. Finally, we use regular weights but randomize the spike timing in the reconstruction step. This is done by randomly permuting, without repetition, the order of time steps of the spike train (the order of time steps is the same for all neurons within the population). For all four types of perturbation, the procedure is repeated *nperm*−times.

#### Decoding with single neurons

We also decode correct choice behavior from the activity of single neurons. We compute the decoding weight with an univariate method, for each neuron independently, and apply the weight to the spike train of the single neuron. We use the same cross-validation procedure as with the multivariate method described above. The univariate decoding weight is computed as the Receiver-Operating Characteristics Curve (*AUC*, [24]), utilizing spike counts. To have univariate decoding weights comparable to multivariate weights, we center the *AUC* score around zero and scale it with the L2 norm,

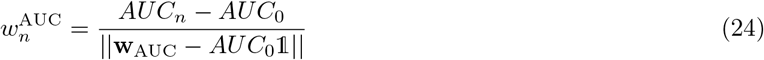

with 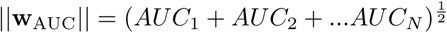, 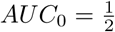 is the at chance prediction and 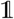 is vector of ones. Univariate decoding weights are then weighting spikes in the hold-out set, for each neuron independently.

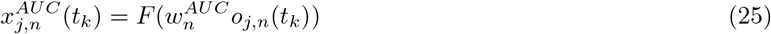

Same as in the multivariate case, we compute the deviation of the signal from the mean,

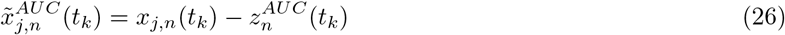

where 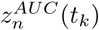 is the average signal across trials from both conditions.

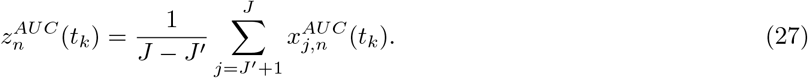

The univariate signal is then averaged across trials, distinguishing conditions “match” and “non-match”.

The null model of reconstruction with single neurons is computed with random permutation of class labels. We compute the AUC score on spike counts with randomly permuted class labels for conditions “match” and “non-match”. Weights are then applied on spike trains, where the label for condition has again been randomly permuted.

#### Correlation analysis

##### Cross-Correlation between the population signals of *plus* and *minus* neurons

We compute the cross-correlation function between population signals of *plus* and *minus* neurons in trial *j*,

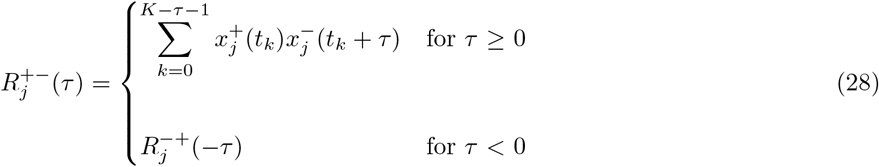

with time lag *τ* = 1, 2, …, 2*K* − 1. The correlation function is normalized with autocorrelation functions at zero time lag,

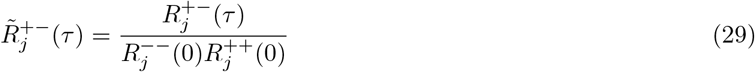

where *R*^++^ (*R*^−−^) is the autocorrelation function for *plus* (*minus*) neurons.

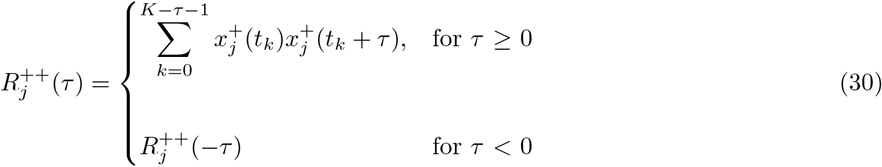

The correlation function is computed in single trials and then averaged across trials, distinguishing conditions “match” and “non-match”,

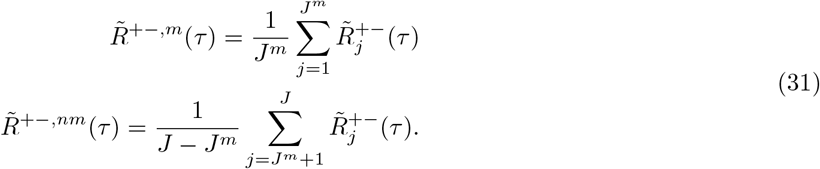

where *J*^*m*^ is the number of trials in condition “match” and trials are ordered such that trials in condition “non-match” follow trials in condition “match”. We estimate the significance of the correlation function with the permutation test. Since there was no difference across conditions, we compute the correlation function using trials from both conditions,

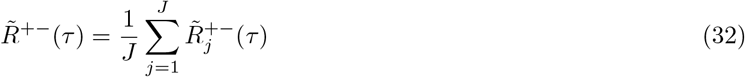

and compare it with the null model. The null model is computed with random weights and random assignment to the group of *plus* and *minus* neurons.

##### Correlation function of the population signals in cortical layers

Similarly, we compute the cross-correlation of population signals between pairs of cortical layers. As before, the correlation function is computed for the two population signals in the same trial,

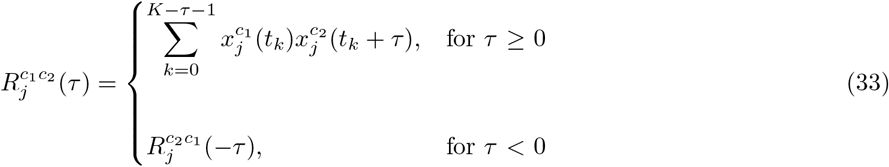

with (*c*_1_, *c*_2_) ∈ {(*SG, G*), (*SG, IG*), (*G, IG*)}. The rest of the procedure is the same as for *plus* and *minus* neurons. The significance of results is evaluated with the permutation test. The null model is computed with random weights and random assignment to one of the three cortical layers.

#### Determining cortical layers from the current source density

The cortical depth can be split into three cortical layers, the superficial or the supragranular layer (SG), the middle or the granular layer (G), and the deep or the infragranular (IG) layer [23]. To split neurons in layers, we designed a method that utilizes patterns of activation of the current source density (*CSD*). *CSD* is a 3-dimensional tensor that associates direction of the current flow to every point in space and time. Normalized *CSD* is defined as *A*_*ijk*_, *i* = 1, …, *N*_*space*_, *j* = 1, …, *N*_*time*_, *k* ∈ [−1, 1], where *i* extends in the spatial dimension, *j* in the temporal dimension and *k* is the direction of the current flow. *CSD* is computed as the second spatial derivative of the trial-averaged local field potential [25]. In general, the G layer is the input layer for sensory stimuli and is characterized by a current sink upon the presentation of a salient visual stimulus, while, simultaneously, the SG and IG layers present a current source [26]. We utilize this pattern of sink and sources to determine the borders of the G layer. First, we search for the strongest current sink in the time window [20,100] ms after the onset of the test stimulus, which is the point in space and time with the maximal value of the current flow, *A*_*max*_(*i′*, *j′*, *k′*). The pattern of sink and sources is primarily a spatial feature and we capture it with the spatial covariance of the current source density,

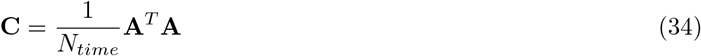

where ^*T*^ denotes the transpose. We now define the vector **c**_*max*_ as the vector of covariance that passes through the point *A*_*max*_.

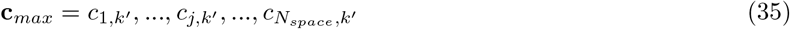

Along this particular vector of covariance, current sinks correspond to peaks and current sources correspond to troughs. We capture the peak that corresponds to the strongest current sink, *c*_*j′*,*k′*_. On each side of this peak, the two troughs correspond to the two current sources of interest. Between the peak and each of the troughs, the vector of covariance crosses the zero line, indicating the current inversion. We determine the upper and the lower border of the G layer as the zero crossing on each side of the peak. After determining the borders of the G layer, units above the upper border of the G layer were assigned to the the SG layer and units recorded below the lower border to the IG layer. In V1, we identified 48, 51 and 61 neurons in the SG, G and IG layer, respectively. In V4, we identified 18 (SG), 42 (G) and 42 (IG) neurons in the respective layers.

## Results

### The population PSTH does not allow to discriminate correct choices on binary stimulus classes “match” and “non-match”

Two adult monkeys *macaca mulatta* have been tested on the visual discrimination task with complex naturalistic images (see methods). In each trial, animal subjects visualized two images, the target and the test, with a delay period in between (Fig 1A). Laminar arrays were inserted into the visual areas V1 and V4, perpendicularly to the cortical surface, spanning the cortical depth (Fig. 1B). We limited the analysis to correct trials and distinguish two conditions, “match’” (correct choice on matching stimuli) and “non-match” (correct choice on non-matching stimuli). Classes “match” and “non-match” are conditioned on a mixed variable that potentially contains the information about both the stimulus and the choice and discriminating the two classes relies on either of these two sources of information or, more likely, a combination of both. Throughout the study, we analyze neural responses in two time windows, corresponding to the interval [0,400] ms with respect to the onset of the target and the test stimuli. The information, necessary for discriminating matching from non-matching stimuli, is only available during the test time window and we therefore expect to discriminate “match” from “non-match” during test, but not during target.

**Fig 1.**
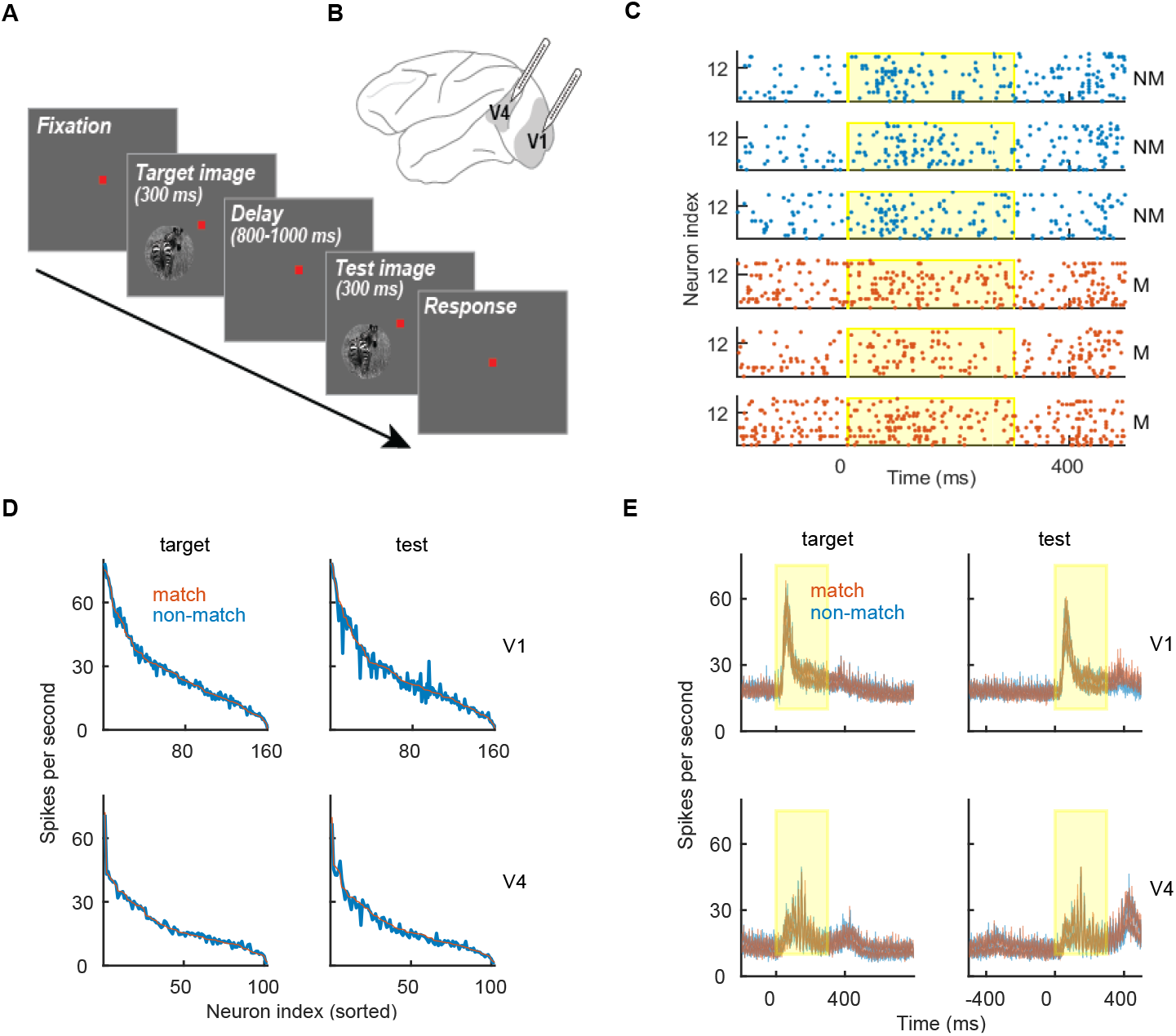
Experimental paradigm and spiking data. **(A)** Experimental paradigm. One trial consisted of the visualization of the target and the test stimuli, interleaved with the delay period. **(B)** Schema of the macaque brain with approximate location of recording sites. **(C)** Spike trains of an example recording session in V4 during the visualization of the test stimulus. We show 3 randomly selected trials in condition “non-match” (blue) and “match” (red). The yellow region marks the presence of the stimulus. **(D)** Mean firing rate of single neurons in V1 (top) and in V4 (bottom) during the target time window (left) and the test time window (right). We show the firing rate in condition “match” (red) and “non-match” (blue). Neurons were collected across all recording sessions and sorted (from strongest to weakest) for the firing rate in condition “match”. **(E)** Population PSTH in conditions “match” (red) and “non-match” (blue). We show the mean ± SEM for the variability across sessions. The presence of the stimulus is marked as the yellow region.

The average firing rate of single neurons is highly variable across neurons but very similar across conditions (Fig 1D). Also, PSTHs in conditions “match” and “non-match” are highly overlapping (Fig 1E). With the population PSTH, the spiking activity is summed across neurons, with each neuron contributing equally to the sum (see methods, eq. 2). In the following, we will assume that the contribution of neurons within the population is not equal, but is weighted according to neuron’s decoding weight. This reflects the fundamental idea that neural networks are not homogeneous ensembles but instead have a structure that allows them to perform computations.

### Weighting spike trains with decoding weights allows to discriminate conditions “match” and “non-match”

We compute decoding weights by training an optimal linear classifier, SVM, on parallel spike counts of simultaneously recorded neurons (see methods). Once decoding weights are learned, they are fixed and can be used in single trials and in real time to compute the population signal. The population signal is computed as a weighted linear sum of spikes (see methods, eq. 12). We collect the population signal across trials, and average it distinguishing conditions “match” and “non-match” (eq. 16). The population signal in conditions “match” and “non-match” is highly overlapping during the target time window (Fig 2A, left). This is expected, since the information, necessary for discrimination, is not yet available. During the test time window, the population signal in conditions “match” and “non-match” diverges (Fig 2A, right), and is significantly different between the two conditions in both V1 and V4 (Fig 2B, right). It can be seen, however, that the temporal profile of the population signal differs across the two brain areas. In V1, the signal diverges early in the trial and stays approximately constant throughout the trial. In V4, the difference between signals in conditions “match” and “non-match” builds up over time and is the biggest towards the end of the trial (Fig 2B, lower right). Interestingly, the difference in signals in V4 slowly oscillates and at the same time increases in every cycle. Considering the population signal as the input current to a read-out neuron, such dynamics would give windows of high and low probability for spiking.

**Fig 2.**
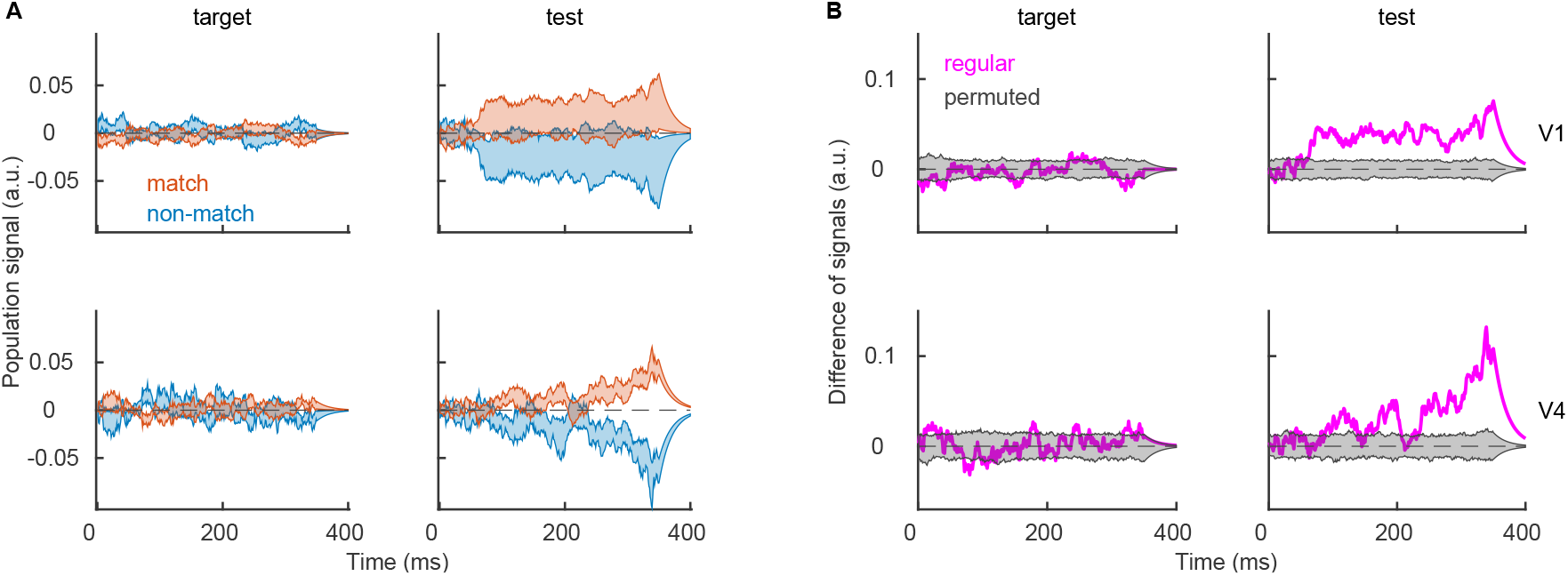
Population signal. **(A)** Population signal in V1 (top) and V4 (bottom) during the target (left) and the test time window (right). Shaded areas indicate the mean ± SEM for the variability across sessions. We show population signals in conditions “match” (red) and “non-match” (blue). **(B)** Difference between session-averaged population signals, showing the regular model (magenta trace) and the distribution of results of the null model (gray area). Parameters: *λ* = 20^−1^ ms, *nperm* = 1000.

The exact time course of the population signal depends on the time constant of the convolution *λ*. With longer time constant, the effect of a spike lasts longer, giving rise to a signal that integrates more over time (Fig 3A, left plots). The use of a linear filter, however, essentially only scales the amplitude of the signal, and this effect can be reversed by rescaling the signal with the area under the temporal filter, Σ_*t*∈*T*_*u*(*t*) (Fig 3A, right plots). As signals are rescaled, the only difference between signals that use different time constants is smoothness, since longer time constants give smoother signal. The oscillatory dynamics of the population signal in V4 therefore cannot be due to the convolution with a particular filter, since the same oscillatory dynamics is present for different time constants (Fig 3A, bottom right) and since it is present for time constants that are shorter than the oscillation cycle. In the rest of the paper, we will use the time constant *λ* = 20^−1^ ms.

**Fig 3.**
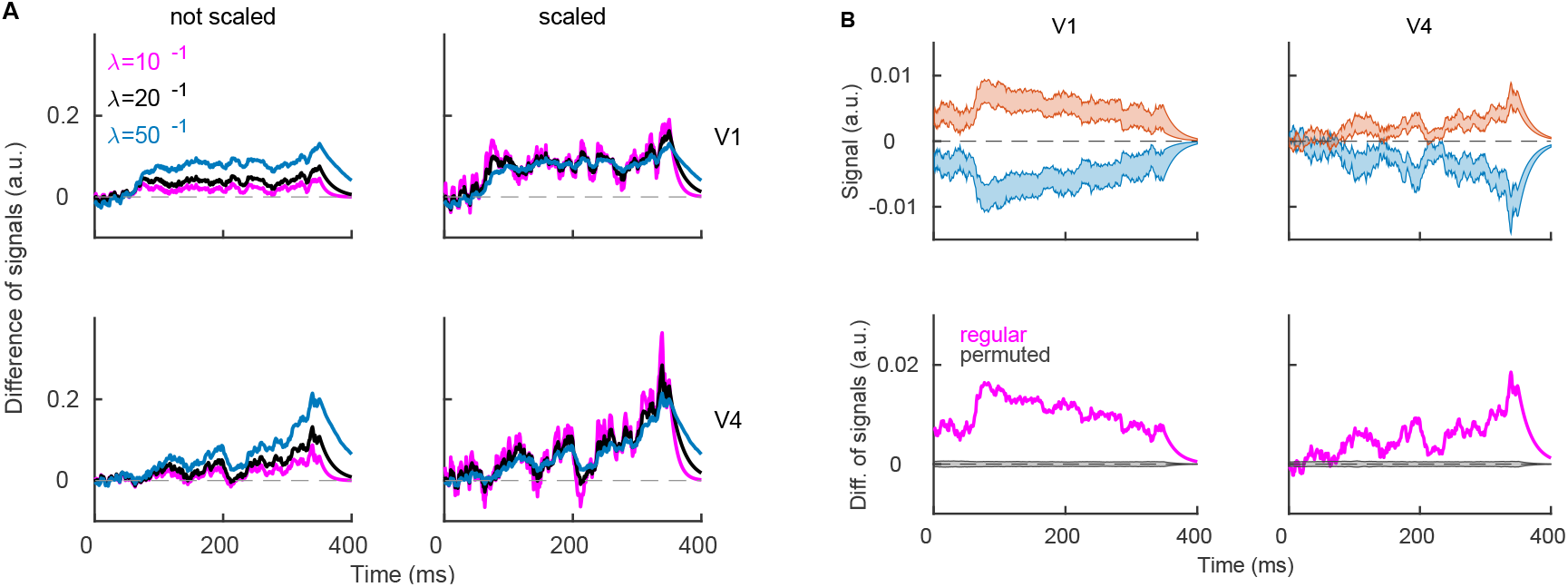
Dependency of the population signal on the time scale of the convolution and on the modulus of weights. **(A)** Difference of population signals during the test time window for different values of parameter *λ*. We use time constants *λ* = [10^−1^, 20^−1^, 50^−1^] ms and plot results without normalization (left) and with normalization (right). **(B)** Decoding with single neurons. Top: Signal, decoded from single neurons in conditions “match” (red) and “non-match” (blue) during the test time window. We show the mean ± SEM for the variability across all recorded neurons in V1 (left) and in V4 (right). Bottom: Difference of signals, averaged across neurons (magenta), and the distribution of the same results from models with permuted class labels (gray). Parameters: *λ* = 20^−1^ ms, *nperm*=1000.

It is also possible to discriminate conditions “match” from “non-match” from the activity of single neurons. We compute decoding weights of single neurons as the area under the ROC curve, and apply the weight of each single neuron to its spike train (see methods, eq. 24 - 27). Results show that the activity of single neurons also allows to discriminate conditions “match” and “non-match”. However, we argue that the univariate approach is less biologically plausible. It is unlikely that any brain structure can isolate the activity of a single synaptic input, while being “bombarded” by a large number of synaptic inputs from hundreds to thousands of neurons [9].

### Correct sign of weights is necessary and sufficient for discrimination

The population signal contains distinct sources of information, and we ask, which of these sources is necessary for discrimination. We disentangle the necessity of a particular source for discrimination by computing the population signal (eq. 16) with a particular source of information being removed (see methods). As we remove the information from the weight vector, population signals in conditions “match” and “non-match” are highly overlapping (p=0.6046 in V1, p=0.8499 in V4, t-test on time-averaged signals with 1000 permutations), indicating that the information in the weight vector is critical for discrimination (Fig 4A, left plots). Next, we disentangle the importance of the sign and of the modulus of weights for discrimination. Removing the information on the sign of weights gives highly overlapping population signals in conditions “match” and “non-match” (p=0.7354 in V1, p=0.4774 in V4, Fig 4, middle left). If, instead, we keep the correct sign and use a random modulus, the population signal in conditions “match” and“non-match” can clearly be discriminated (p*<* 10^−8^ in both areas, Fig 4A, middle right). Finally, as we remove the information in spike timing, discrimination remains possible (p*<* 10^−8^ in both areas, Fig 4A, right plots), showing that the spike timing is not critical for discrimination. However, permutation of spike timing removes the temporal profile of the population signal, including the oscillating dynamics in V4. Nevertheless, we conclude that the information contained in the sign of weights is necessary and sufficient for discrimination of conditions “match” and “non-match”.

**Fig 4.**
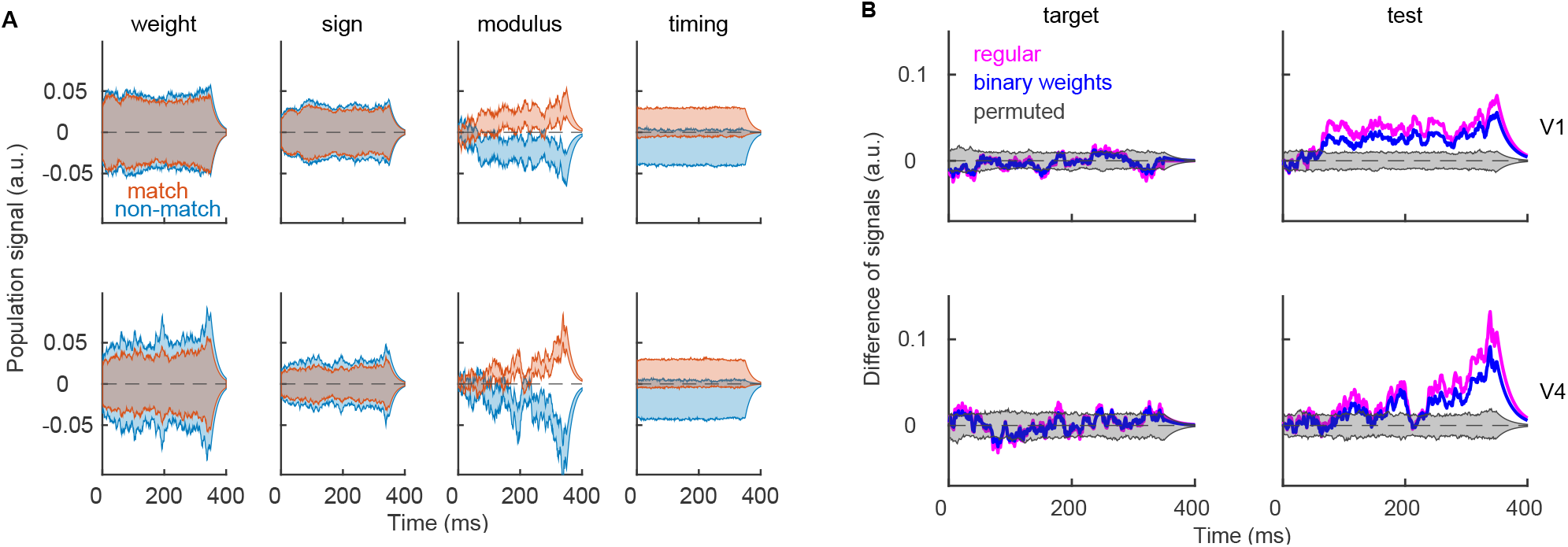
Distribution of population signals for models with specific permutation. **(A)** Population signal during the test time window, computed with random weights (left), random sign of weights (middle left), random modulus of weights (middle right) and randomly permuted spike timing (right). We show results in V1 (top) and in V4 (bottom) in conditions “non-match” (blue) and “match” (red). For all procedures, we used 1000 random permutations and we plot the entire distribution. **(B)** Difference of population signals for the regular model (magenta) and for the model with binary weights (blue). The gray area marks the distribution of results using models with permuted class labels. Parameters: *λ* = 20^−1^ ms, *nperm* = 1000.

Results have shown that the modulus of weights is not necessary for discrimination, and we compute the population signal using binary weights to see if such a population signal is similar to the regular population signal (as on Fig 2). All positive weights are set to the same value, *a*, and all negative weights are set to its inverse, −*a*. We estimate the scalar *a* in such a way that the range of the weight vector is the same as with original set of weights, i.e., 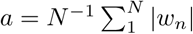. The population signal with binary weights is indeed similar to the population signal with regular weights, and allows discrimination of conditions “match” from “non-match” (Fig 4B). Optimal (and heterogeneous) weights still have a slightly greater discriminatory power than binary weights (e.g., the difference of signals in “match” and “non-match” is greater with optimal weights). Nevertheless, since the loss of discriminatory power is relatively small, we can relax the hypothesis of optimality, questionable in the biological network, to the correct assignment of the sign of weights.

### Neurons with positive and negative weights respond with anti-symmetry

Since the sign of weights is the crucial source of information for the model, we ask, how do *plus* and *minus* neurons contribute to the population signal. By design, *plus* and *minus* neurons have the opposite effect on classification. The weight vector determines the direction of the plane that separates data points, and *plus* and *minus* neurons rotate the separating plane in the opposite directions. We therefore split neurons in two subpopulations according to the sign of weights and compute the population signal with each of the two subpopulations (see methods, eq. 18). During test, neurons with positive and negative weights respond with anti-symmetry (Fig 5A). *Minus* neurons increase the activity above the baseline in condition “non-match” and decrease below the baseline in condition “match”, while *plus* neurons do the opposite. The divergence of the signals in the two conditions most prominently appears at the end of the trial, which could be due to a neuron-type specific feedback from higher brain areas. In V4, the anti-symmetric response is observed only with *plus* neurons, while *minus* neurons do not have any significant discriminatory capacity (Fig 5A, bottom). During the target time window, as expected, the population signal fluctuates around zero in all cases (Supplementary Fig S1). We conclude that *plus* and *minus* neurons during test respond with anti-symmetry, with the exception of *minus* neurons in V4, that do not seem to discriminate conditions “match” from “non-match”.

**Fig 5.**
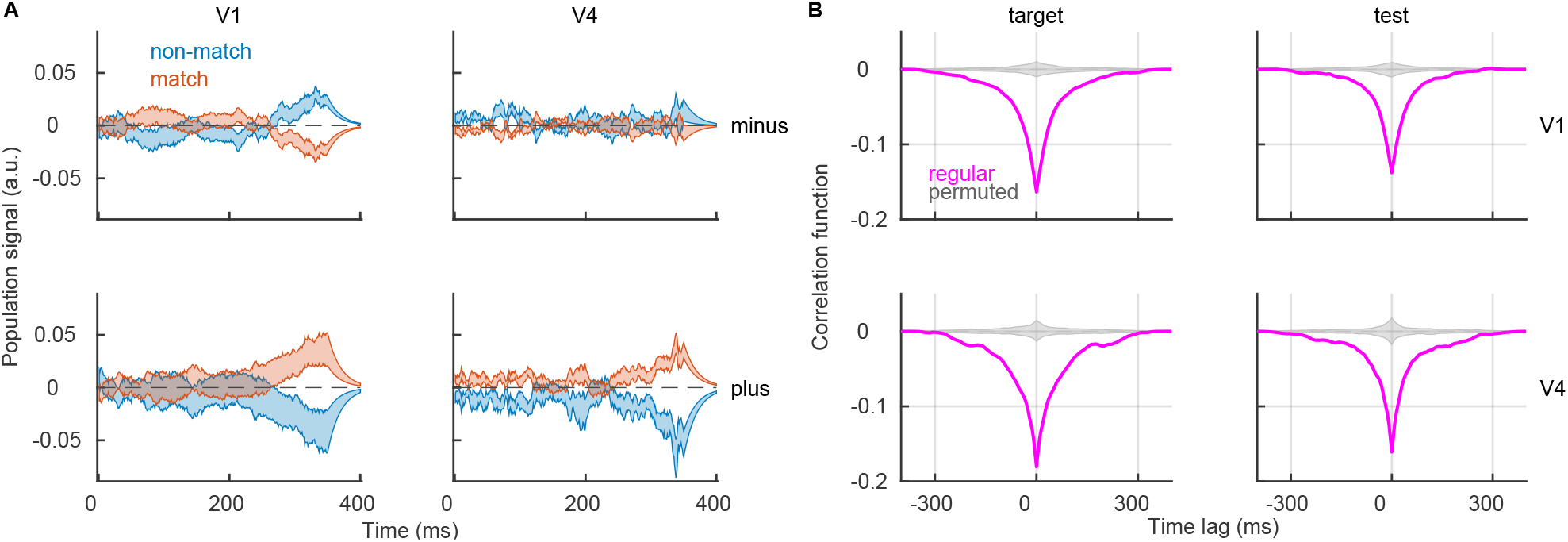
*Plus* and *minus* neurons respond with anti-symmetry. **(A)** Population signal of *minus* neurons (top) and *plus* neurons (bottom) during the test time window. We show the mean ± SEM for the variability across recording sessions and report results in condition “match” (red) and “non-match” (blue). **(B)** Cross-correlation function between population signals of *plus* and *minus* neurons. To compute the cross-correlation function, we use trials from both conditions. Results are shown for the true model (magenta) and for models with random assignment to the subpopulation of *plus* and *minus* neurons. Parameters: *λ* = 20^−1^ ms, *nperm* = 1000.

The low-dimensional signal for *plus* and *minus* neurons is computed by splitting the population of simultaneously recorded neurons into two subpopulations. The signals of *plus* and *minus* neurons therefore evolve simultaneously within the same trial. We compute the interaction between the two signals with the cross-correlation function (see methods, eq. 28-32). Interestingly, there is a negative correlation between the population signals of *plus* and *minus* neurons, in the two brain areas and in both target and test time window (Fig 5B). There is no significant difference between correlation functions in conditions “match” and “non-match” (Supplementary fig S1 D). Negative interaction between *plus* and *minus* neurons appears as a strong and robust effect in both brain areas.

### The superficial layer of the cortex discriminates best conditions “match” and “non-match”

In the previous section, neural populations were split according to the sign of decoding weights. In the last part, we split the neural population according to the spatial location of neurons across the cortical depth in three cortical layers (superficial or SG, middle or G and deep or IG layer, see methods). We split neurons in layers using a method based on the spatial covariance of the current source density (Fig 6, see methods). The population signal is computed in each layer separately (eq. 23). Layer-specific population signals reveal that the superficial layers have the strongest discriminatory capacity of conditions “match” and “non-match” in both V1 and V4 (Fig 7). In V1, the middle layer shows a weak effect during the first half of the trial (Fig 7B, middle left) and in V4, middle and deep layers show a weak effect towards the end of the trial (Fig 7B, middle and bottom right). This is true for the test time window, while during the target time window, the population signal in all layers stays close to zero (Supplementary Fig S2). The deep layer of V1 shows almost no discriminatory capacity in either test (Fig 7A, B, bottom left), or target (Supplementary Fig S2).

**Fig 6.**
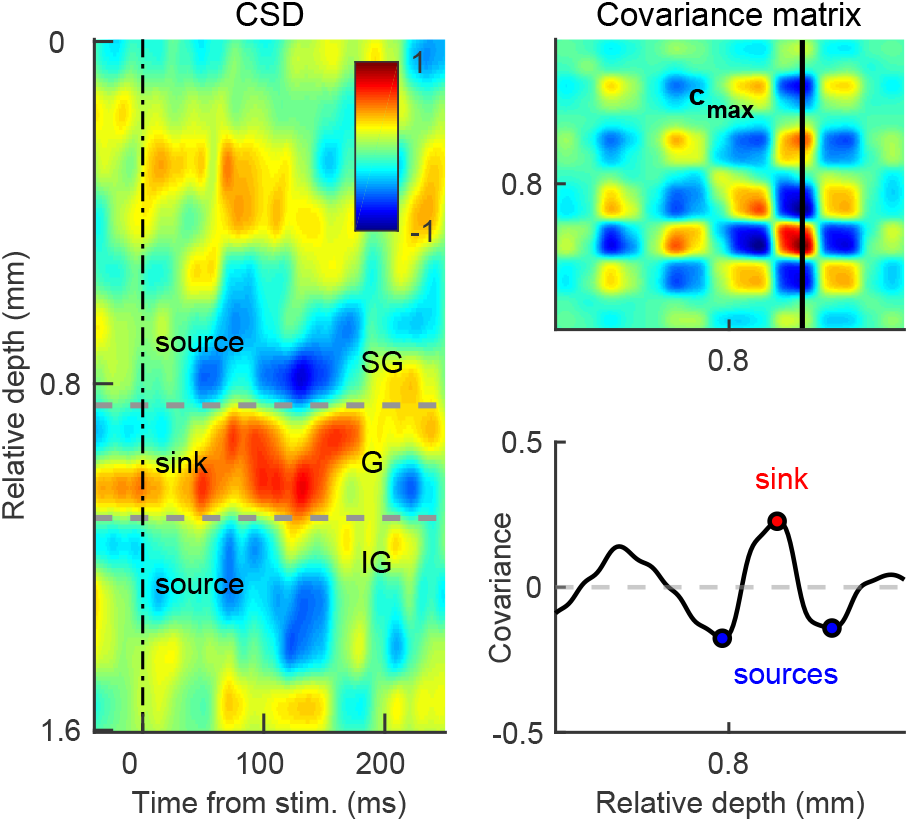
Method for assignment of cortical layers from the current source density. Left: Current source density (*CSD*) during one recording session in V4. The x-axis shows time, relative to the onset of the test stimulus, and the y-axis shows the cortical depth, relative to the position of the upper channel on the laminar probe. The color indicates the direction of the current flow, with current sinks in red and with current sources in blue. After the onset of the stimulus, we observe a characteristic pattern of current sink and sources. The sink is a hallmark of the granular (G) layer, while sources characterize the supragranular (SG) and the infragranular (IG) layers. Top right: The pattern of sinks and sources is captured by the spatial covariance of the current source density. We select the vector of covariance that passes through the strongest sink, **c**_*max*_. Bottom right: Plotting the vector of covariance as a function of the cortical depth, one of the peaks corresponds to the strongest current sink (red point) and neighboring troughs correspond to current sources (blue).

**Fig 7.**
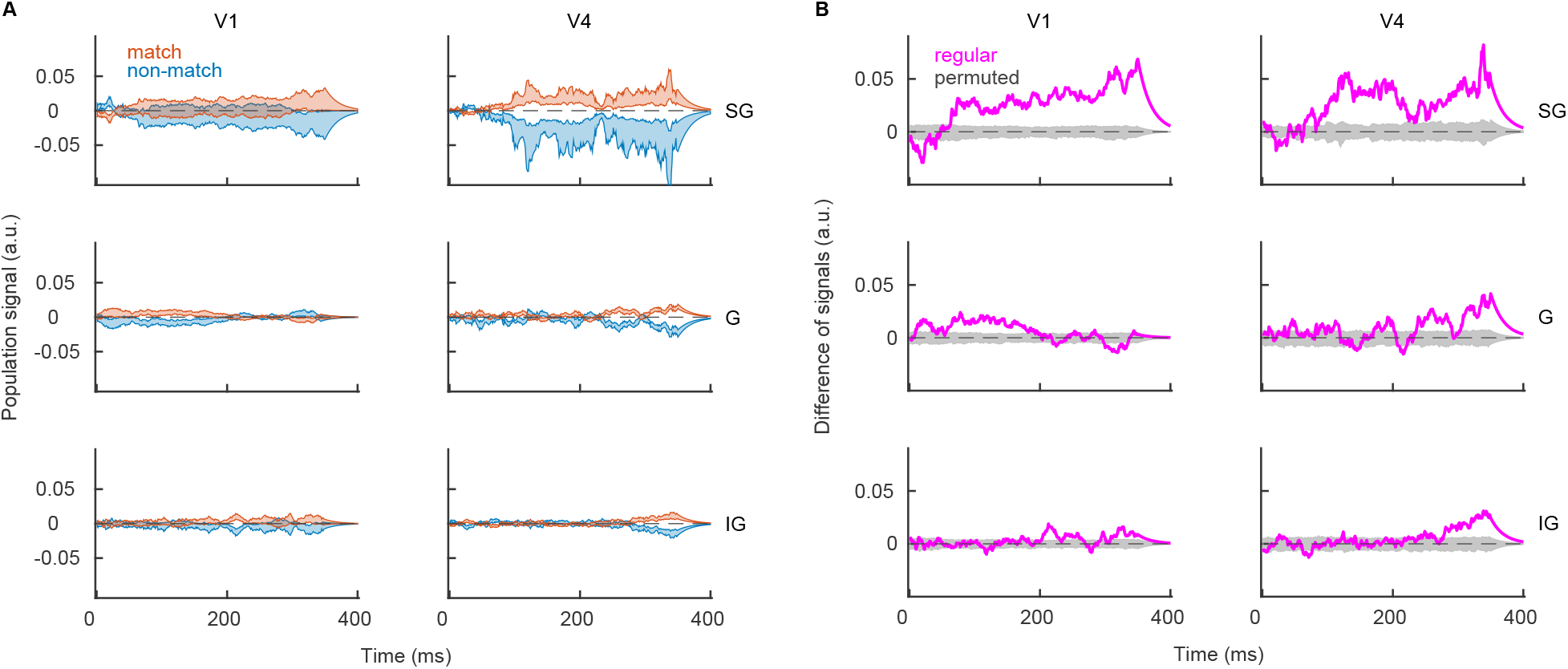
The superficial layer of the cortex is the best in discriminating conditions “match” from “non-match”. **(A)** Population signal in recording sessions in the superficial (top), middle (middle) and deep cortical layers (bottom) in V1 (left) and in V4 (right). We show the mean ± SEM across recording sessions in conditions “match” (red) and “non-match” (blue). **(B)** Same as in **A**, but for the session averaged difference of signals. We show results of the regular model (magenta) and the distribution of results for the model with permutation (gray). Parameters: *λ* = 20^−1^ ms, *nperm* = 1000.

With layer-specific reconstruction of spike trains, we obtain three simultaneous population signals, one in each layer. We measure the linear correlation of population signals for each pair of layers with the cross-correlation function (see methods, eq. 33). Results show positive correlation across all pairs of layers during both test (Fig 8) and target time window (Supplementary Fig S3 A). There is no significant difference across conditions “match” and “non-match” in neither target nor test (Supplementary Fig S3 B,C). We conclude that positive correlation across layers is a robust and generic property of the cortex, similarly to the negative correlation between subnetworks of *plus* and *minus* neurons.

**Fig 8.**
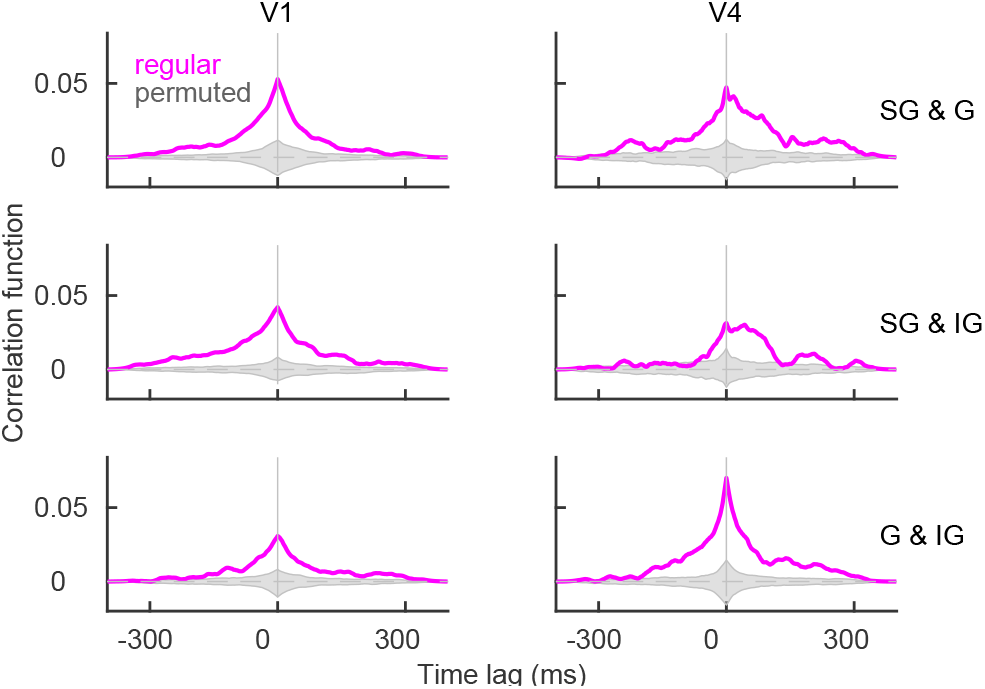
Population signals are positively correlated across the cortical layers. The cross-correlation function for pairs of cortical layers, SG & G (top), SG & IG (middle) and G & IG (bottom) during the test time window. The cross-correlation function uses trials from both conditions. We plot results of the regular model (magenta) and distribution of results for models with permutation (gray). Parameters: *λ* = 20^−1^ ms, *nperm* = 1000.

## Conclusion

We presented a new model of the read-out of parallel spike trains that exploits the structure of the population code. We assumed the point of view of a read-out neuron, receiving synaptic inputs from a population of projecting neurons. The gist of the present read-out method is to endow a biological neural network with functionality. This is done by assuming that decoding weights between projecting neurons and a hypothetical read-out unit are such as to allow classification. Decoding weight determines how does the spike of a particular neuron affect the population signal. After weighting spikes and summing across neurons, we get a 1-dimensional signal, evolving in real time, that can be thought of as the sum of post-synaptic potentials at the read-out unit, modulating the spiking probability of the latter. We have shown that the population signal clearly allows to discriminate conditions “match” and “non-match” in V1 and V4 during the test time window, when we would expect it to do so. Importantly, the population signal does not allow the discrimination during the target time window, when we would not expect it to do so. We demonstrated that discrimination critically depends on the correct assignment of the sign of the weight. Results show that neurons with the opposite sign of weight respond in anti-symmetric fashion to the mutually exclusive stimulus classes “match” and “non-match” and that population signals of neurons from the opposing coding pools are negatively correlated (for a related analysis in the retina, see [27]). Distinguishing superficial, middle and deep layers of the cortex, we show that the superficial layer is the most important in discriminating the two behavioral conditions in both brain areas.

Present model gives insights in the structure of the population code and into how this structure allows computation at the level of neural networks. The model can be applied to any data set with parallel spike trains where it is possible to assume the nature of the computation that underlies the neural activity. While the concept of weighted spike trains is generally applicable, the way the population decoding weights are computed has to be adapted to the specific case at hand. Here, we computed decoding weights using a specific supervised learning method (linear SVM), but another supervised learning method, unsupervised learning or statistical method can be used instead. Our choice of the linear SVM is justified with SVM’s optimality for binary classification tasks and we would expect that the use of another classifier would results in a decoding model with decreased performance. We argue that, in the present context, supervised learning is a plausible assumption. The animal is rewarded for the correct behavior (which is the one we analyze here) and the reward signal could enact the teaching signal, assumed by supervised learning. Moreover, we also show that the optimality assumption can be relaxed with only a small loss of predictive power.

In the present experimental setting, the behavioral task consists in matching delayed samples. The decision of the animal (“same” or “different”) is based on the comparison of the test stimulus with the stimulus from the past (the target stimulus). This is a relatively complex cognitive task that presumably requires the activation of the working memory [28]. Visual areas, in particular their superficial layers, receive top-down projections [29], while, at the same time, they are also driven by the bottom-up inputs. It has been shown theoretically that in the presence of the common top-down input, population-wise coding weights can be learned with local synaptic plasticity rules [17]. We speculate that after learning, the top-down input to V1 and V4 could selectively target *plus* or *minus* neurons, depending on the condition. Such a context-dependent top-down signal could be computed in the prefrontal cortex, as described in [30]. Neurons with positive weight would be preferentially targeted in condition “match” and neurons with negative weight in condition “non-match”, and such a condition-specific top-down signal could explain the anti-symmetric activation of *plus* and *minus* neurons that we observe towards the end of trial. Among the three cortical layers that have been tested, the superficial layers have shown the best discrimination capacity. The strength of the effect in the superficial layers of V1 and V4 with respect to other layers corroborates our idea of the top-down influence on representation, since the superficial layers receive top-down inputs most abundantly and are as such best suited to perform the required computation.

From the biophysical point of view, it has been understood that variable spiking of single neurons can be mechanistically accounted for by the high-conductance regime [31], where the stream of excitatory and inhibitory synaptic inputs largely cancel each other out [32]. While the mean excitatory and inhibitory currents cancel each other out, remaining fluctuation of the membrane potential gives rise to variable spiking. It has been proposed that similar mechanism operates at the level of the population signal, where neurons with opposite coding function cancel out each other’s effect [15], and that such canceling is critical for the correct representation of the population signal. In this respect, the method presented here is fundamentally different from methods that do not assume a coding function and where the dimensionality reduction consists in collapsing the dimensionality of homogeneous neural ensembles (with a population PSTH, as done here, or using a more advanced method). According to our results, it is possible to simplify the read-out of cortical spike trains by assuming binary read-out weights (e.g., for the purpose of the analytical treatment). However, any further simplification would take away the ability of the neural ensemble to perform computation. It is interesting to consider the striking difference between the population PSTH and the population signal during the target time window. While we observe a strong peak of activity after the stimulus onset with the former, the latter stays around zero. Presumably, this is so because neurons with opposite coding function cancel out each other’s effect and maintain the representation of the “zero” signal, as suggested in [15].

Comparing our model with Generalized Linear models and other models of Stimulus-Response Functions (SRF, see [33] for a review), the two have in common the goal of the analysis - modeling neural activity from the functional perspective. However, present read-out model importantly differs from models of SRF in several ways. First, models of SRF are encoding models while ours is a decoding model. Second, SRFs model the response function of single neurons and implicitly assume that the representation of the stimuli can be captured by spatio-temporal filters of single neurons. Our model, on the contrary, assumes that task variables are represented by a distributed code and encoded jointly by the entire network. Third, while models of SRF cannot be mapped on low-level physical properties of biological networks, our model presumably can. Potentially, coding with filters and coding with distributed codes might be two complementary rather than competing approaches that the brain utilizes for processing of visual stimuli. While coding with filters might be advantageous with simplistic, low-dimensional stimuli (e.g., moving bars, gratings, moving dots, etc…), distributed population code could be used by the brain to process complex natural stimuli in a more high-level and ecological setting.

In the present work, we have assumed, for simplicity, that all neurons within the population project to the same read-out unit. We argue that in our case, this assumption is reasonable, since neural populations have been recorded across the cortical depth. Because of the retinotopic organization of the visual cortex [34, 35], neurons that span the cortical depth perpendicularly to the surface share a large proportion of their inputs, and project, at least partially, to the same read-out units. Moreover, it has to be emphasized that removing some of the neurons from the population does not change much the population signal, as long as the sign of weights of remaining neurons is correctly assigned. Nevertheless, it would be interesting to directly verify the validity of present results with an experimental assay in the behaving animal, where the activity projecting neurons and of the read-out neuron is monitored simultaneously, the experiment envisioned by Hubel and Wiesel [1]. Recordings from behaving animal still give only a small subsample of all the units that are active in parallel in biological networks. On the modeling side, an interesting way of extending present work would consist in simulating a realistic model of the cortical column and endow it with coding functionality, as it follows from present analysis. Having a realistic number of neurons and biologically plausible connectivity structure would allow to estimate how does the discrimination capacity of the network behave with bigger number of neurons, the presence of excitatory and inhibitory neurons, etc. Such a data-driven model could be studied analytically with advanced methods of dimensionality reduction [36, 37] that provide an accurate description of the population firing rate.

## Supporting information

S1 Fig. Sign-specific population signal.

S2 Fig. Population signal in cortical layers during the target time window.

S3 Fig. Correlation function across cortical layers.

## Acknowledgments

We thank Caroline Matthis and Robert Seidl for comments on previous draft.

