## Supplementary figure 1 for "Reading-out task variables as a low-dimensional reconstruction of neural spike trains in single trials"

### S1 Fig: Sign-specific population signal.

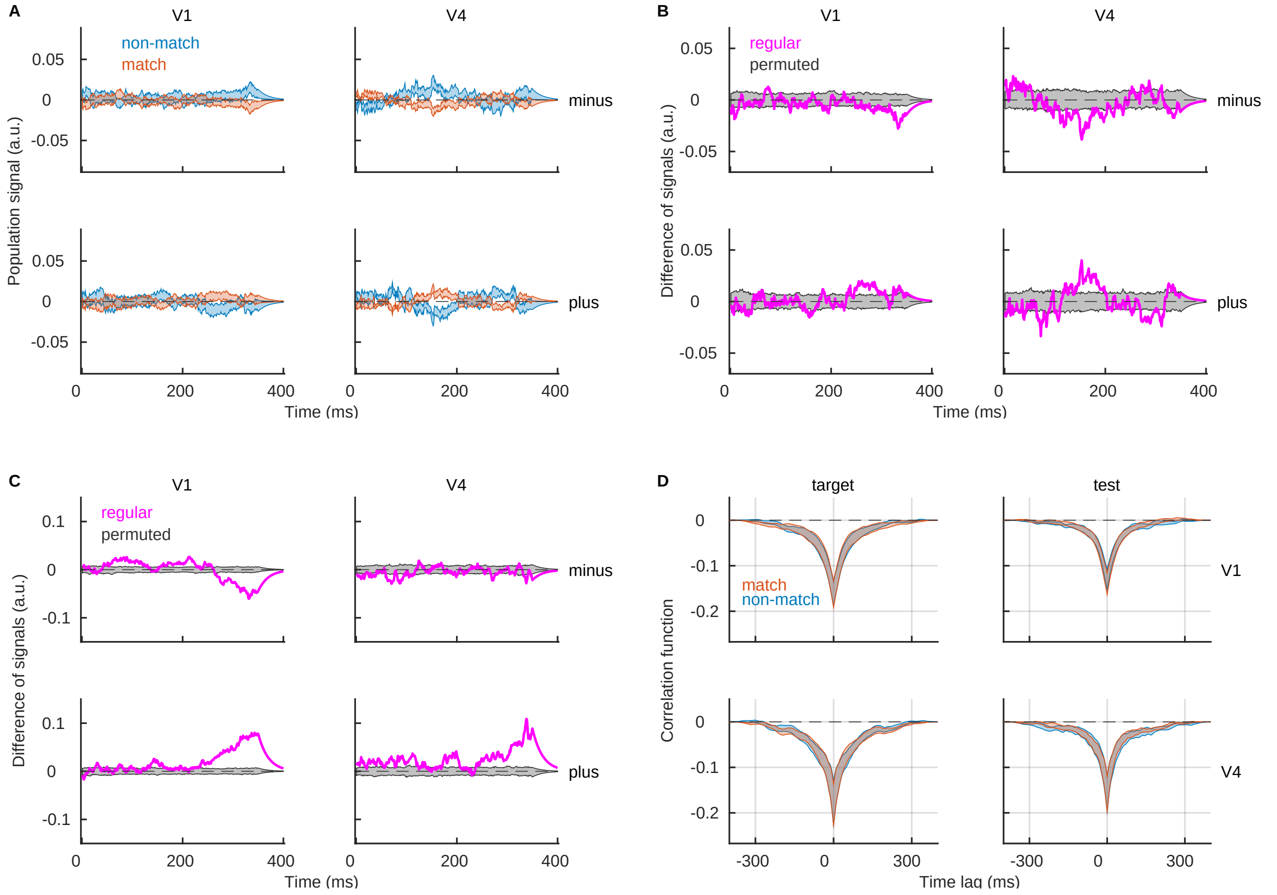


**(A)** Population signal for minus neurons (top) and plus neurons (bottom) during target time window. We show the mean ± SEM across recording sessions in conditions "match" (red) and "non-match" (blue). **(B)** Session-averaged difference of population signals during target. We show the result of the regular model (magenta) and the distribution of results of models with permuted class labels (gray). **(B)** Same as in A, but for the test time window. **(D)** Correlation function between population signals of plus and minus subpopulations in condition "match" (red) and "non-match" (blue). Parameters: $\lambda = 20^{-1}\ ms$, $nperm = 1000$.
