## Supplementary figure 2 for "Reading-out task variables as a low-dimensional reconstruction of neural spike trains in single trials"

### S2 Fig: Population signal in cortical layers during the target time window.

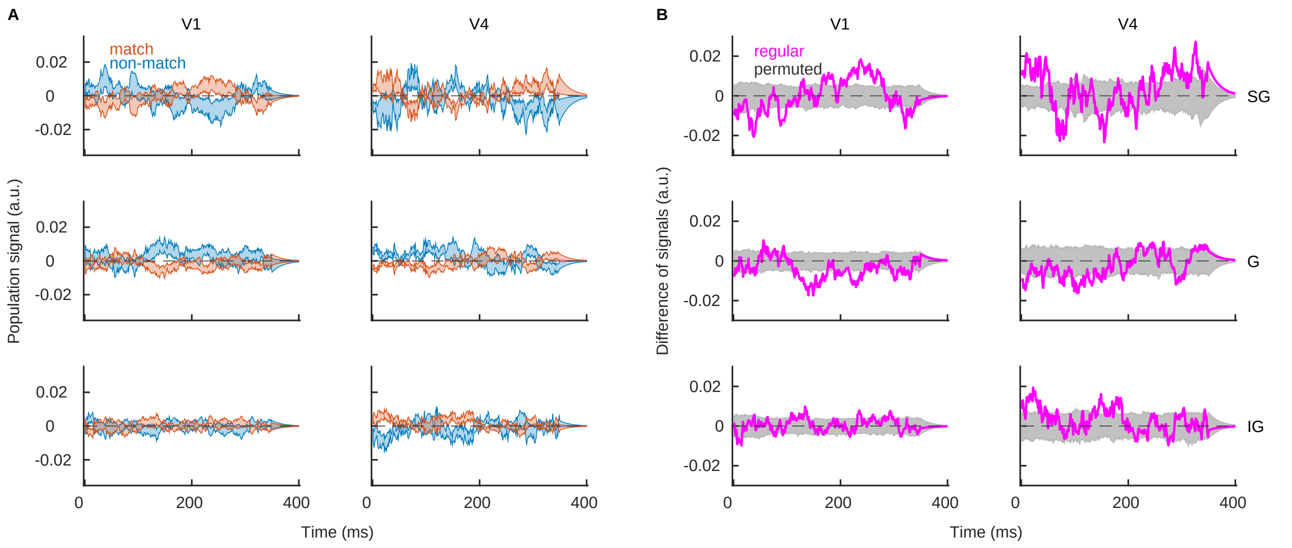


**(A)** Population signal in the superficial (top), middle (middle) and deep cortical layers (bottom) in conditions "match" (red) and "non-match" (blue). We show the mean ± SEM in V1 (left) and in V4 (right), where SEM is for the variability across recording sessions . **(B)** Same as in **(A)**, but for the session averaged difference of signals, $x^{diff} = \tilde{x}^m - \tilde{x}^{nm}$. We show the result of the regular model (magenta) and the distribution of results for the model with permuted class labels (gray). Parameters: $\lambda = 20^{-1}$ $ms$, $nperm = 1000$.
