## Supplementary figure 3 for "Reading-out task variables as a low-dimensional reconstruction of neural spike trains in single trials"

### S3 Fig: Correlation function across cortical layers.

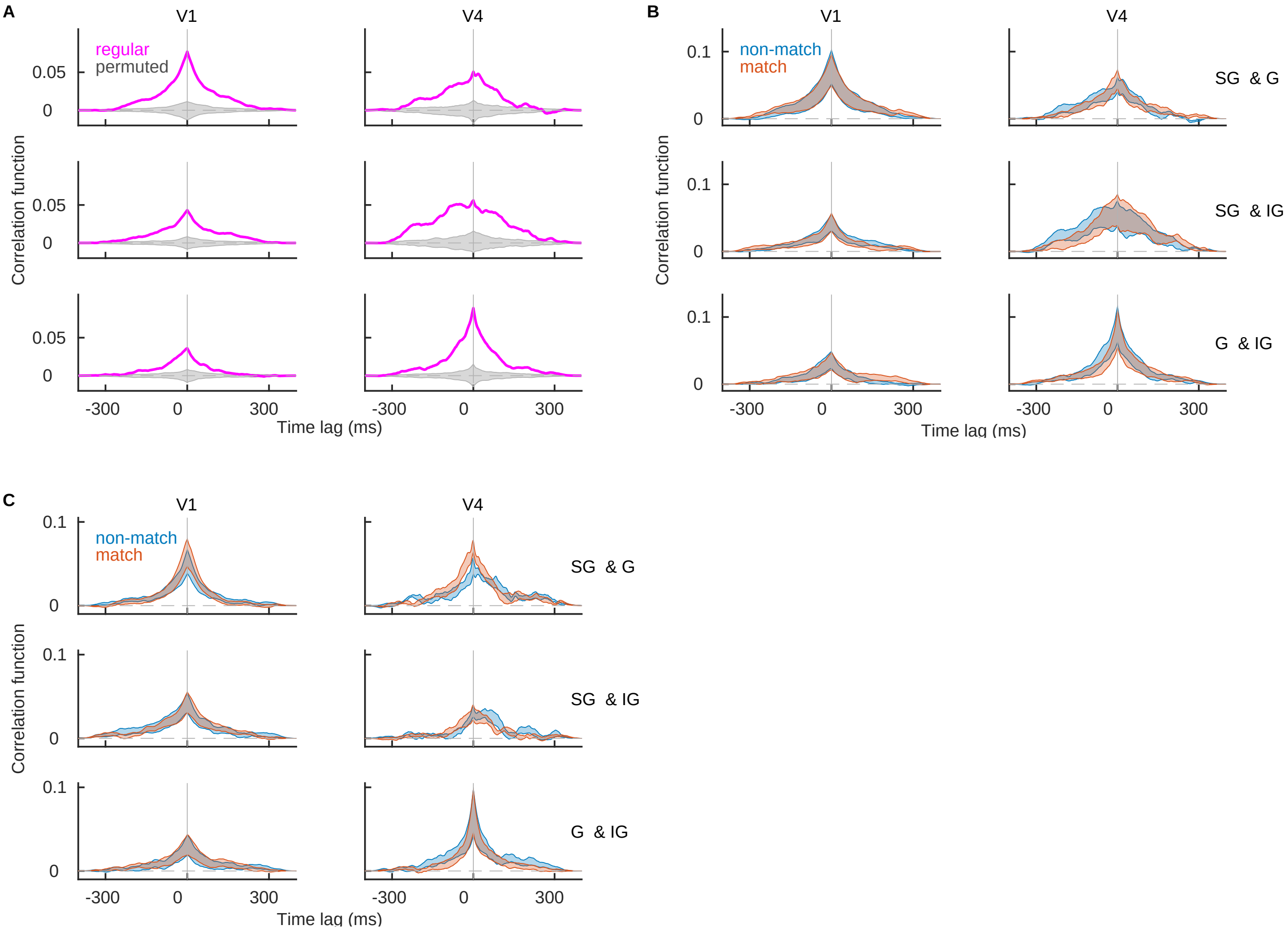


**(A)** Correlation function across pairs of cortical layers SG & G (top), SG & IG (middle) and G & IG (bottom) during the target time window. Correlation function is computed from trials of both conditions. The magenta trace shows the result of the regular model and the gray region shows the distribution of results of models with permutation (the entire distribution). **(B)** Same as in **(A)**, but showing results separately for condition "match" (red) and "non-match" (blue). **(C)** Same as in **(B)**, but in test time window. Parameters: $\lambda = 20^{-1}\ ms$, $nperm = 1000$.
